# Deep learning models to map osteocyte networks can successfully distinguish between young and aged bone

**DOI:** 10.1101/2023.12.20.572567

**Authors:** Simon D. Vetter, Charles A. Schurman, Tamara Alliston, Gregory G. Slabaugh, Stefaan W. Verbruggen

## Abstract

Osteocytes, the most abundant and mechanosensitive cells in bone tissue, play a pivotal role in bone homeostasis and mechano-responsiveness, orchestrating the intricate balance between bone formation and resorption under daily activity. Studying osteocyte connectivity and understanding their intricate arrangement within the lacunar canalicular network (LCN) is essential for unraveling bone physiology. This is particularly true as our bones age, which is associated with decreased integrity of the osteocyte network, disrupted mass transport, and lower sensitivity to the mechanical stimuli that allow the skeleton to adapt to changing demands. Much work has been carried out to investigate this relationship, often involving high resolution microscopy of discrete fragments of this network, alongside advanced computational modelling of individual cells. However, traditional methods of segmenting and measuring osteocyte connectomics are time-consuming and labour-intensive, often hindered by human subjectivity and limited throughput. In this study, we explore the application of deep learning and computer vision techniques to automate the segmentation and measurement of osteocyte connectomics, enabling more efficient and accurate analysis. We compare several state-of-the-art computer vision models (U-Nets and Vision Transformers) to successfully segment the LCN, finding that an Attention U-Net model can accurately segment and measure 81.8% of osteocytes and 42.1% of dendritic processes, when compared to manual labelling. While further development is required, we demonstrate that this degree of accuracy is already sufficient to distinguish between bones of young (2 month old) and aged (36 month old) mice, as well as capturing the degeneration induced by genetic modification of osteocytes. By harnessing the power of these advanced technologies, further developments can unravel the complexities of osteocyte networks in unprecedented detail, revolutionising our understanding of bone health and disease.

## Introduction

Osteocytes are the mechanosensitive cell underlying the exquisite ability of bone to finely balance formation and resorption in response to daily activities^1^. These cells are spread throughout bone tissue, residing within a vast system of lacunae and canaliculi. This system, known as the lacunar-canalicular network (LCN), forms a network of intricately connected architecture that is crucial for maintaining nutrient supply for osteocytes distant from the vasculature^2,3^. Within the LCN the osteocyte processes biochemical signals that regulate bone matrix production^4,5^, mineralisation^6^ and resorption^7,8^ such that skeletal strength is maintained^3^. In this manner, the osteocyte network contributes to calcium homeostasis, but also regulates phosphate homeostasis at the kidney via release of FGF23^9^. This network is now considered to act as a vast endocrine organ, as it is proposed to regulate myelopoiesis^9^, as well as glucose metabolism^10^ and fertility^11^ via osteocalcin secretion.

By far the most abundant bone cell type, osteocytes act as a network of strain gauges, with the connections between them allowing the integration of signals to the osteoblasts and osteoclasts at the bone surface^12^. Furthermore, as osteocytes are highly mechanosensitive, the local parameters of the LCN immediately surrounding individual osteocytes can affect their mechanosensitive response^13–17^ and, with growing evidence that osteocytes have the ability to alter this local geometry themselves^18–20^, it is also possible that they may tune their local environment to maintain homeostatic stimulation^21^. Perhaps most indicative of the importance of this osteocyte network, recent studies have found significant disruption to this system occurs alongside broader degeneration of properties in aged bone, with degradation of the lacunar-canalicular network (LCN)^22^, including loss of canaliculi and changes in lacunar geometries^23–25^. However, most studies of the LCN to date are confined to small field of view within microscopy sections of a femur or tibia, with computational modelling mostly focusing on the mechanical stimulation of single cells. While connectomics techniques do indeed exist in other fields (e.g. neurobiology) and are beginning to be applied to the LCN, to date they are similarly confined to small regions and imaging datasets. Since robust quantitative analysis of the LCN is critical to understanding the role and regulation of osteocytes in bone health, aging, and disease, the continued reliance on time-consuming manual segmentation remains a limiting factor for the field. Therefore, what is clearly required is a rapid and automated analysis tool to segment and measure osteocyte connectomics, in disease and health.

The significant recent advances in machine learning models in biomedical imaging present a promising opportunity for osteocyte researchers. Deep learning models, such as Convolutional Neural Networks (CNNs), are now the foundation of numerous high-performing medical image segmentation models^26–29^. In particular, U-Net, developed by Ronneberger et al.^30^, has gained widespread acceptance in biomedical image segmentation due to its ability to generalise across diverse tasks and perform effectively even with limited amounts of labelled data. However, due to the limited kernel size and inherent locality of the convolution operation, these CNN-based approaches have exhibited limitations in learning long-range dependencies and global context, which are vital for segmenting complex datasets^31,32^. The recent successes of Transformers in the Natural Language Processing (NLP)^33^ field has inspired researchers to adapt this novel method to the vision domain. These models employ self-attention mechanisms to construct comprehensive encoder-decoder architectures, facilitating the capture of long-range dependencies inherent in image data. Numerous studies have integrated the transformer into medical image segmentation^34–36^, exhibiting results that either match or exceed the current state-of-the-art. Additionally, the Swin-UNet model leverages the adaptability and self-attention benefits of transformers along with the localised feature extraction capabilities intrinsic to the U-Net design^31^. This hybrid approach showcases the potential of integrating the strengths of different architectural designs to improve the performance of image segmentation tasks. Nevertheless, image segmentation remains a formidable challenge. Particularly, in the context of the LCN, semantic segmentation encounters several unique obstacles. These include image noise from microscope images, varying illumination and staining conditions in cellular microscopy (due to irregularities of dye absorption), irregularities in cell shapes, and the presence of abnormalities within the data (Figure 1).

**Fig. 1:**
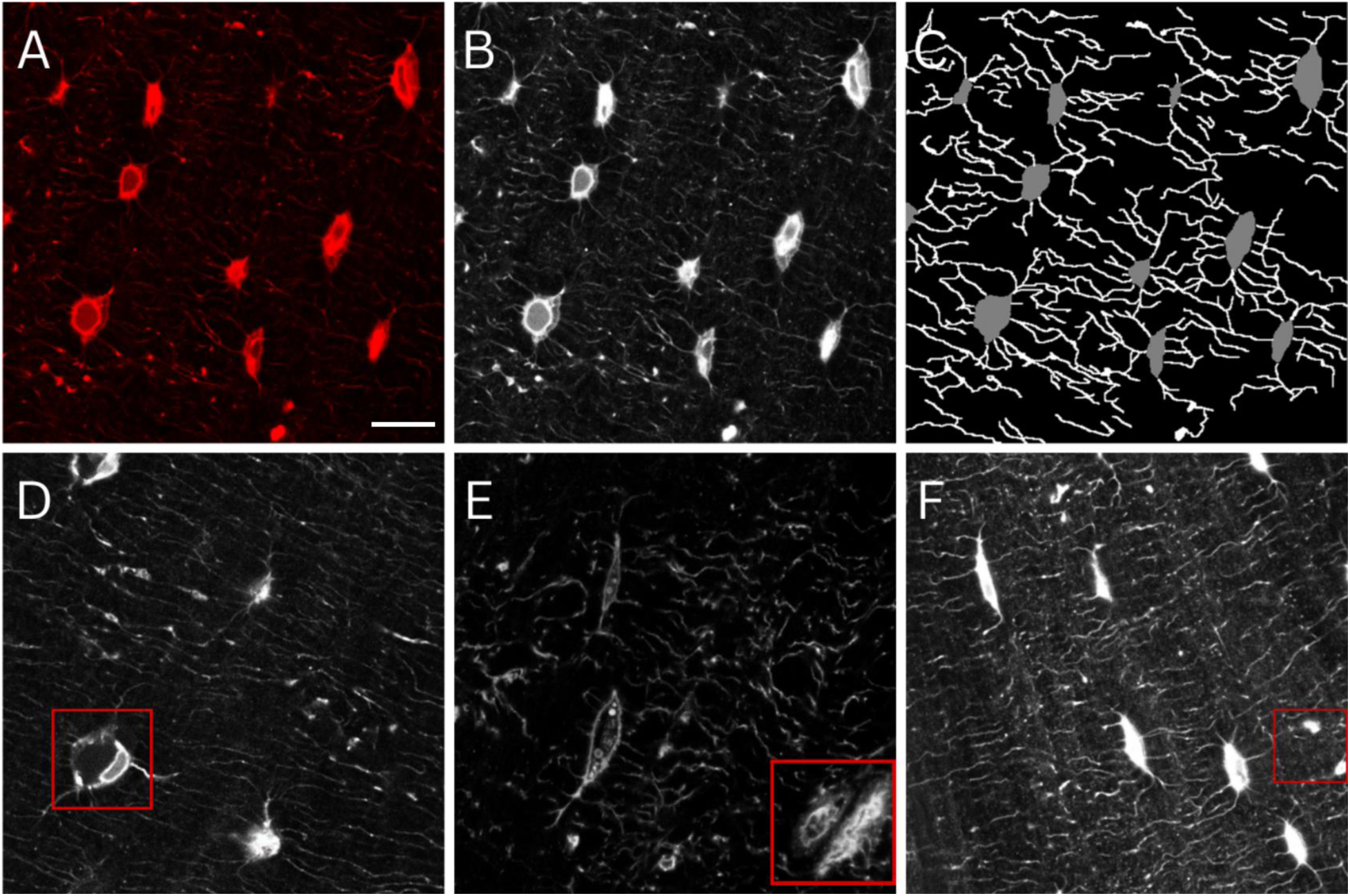
Osteocytes and dendrites manually labelled to train deep learning models. (A) Confocal laser scanning microscopy images of osteocytes (red, phalloidin, cytoskeleton). (B) Images were greyscaled for training, (C) followed by manual annotation and labelling of osteocytes and dendrites. Raw images contained features that presented significant challenges to deep learning models, including (D) shrunken osteocytes possibly fixed during apoptosis, (E) transiting blood vessels, and (F) locations of fused dendrites or the edge of an out-of-plane osteocyte. Scale bar = 10 µm.

Perhaps due to these challenges, to date, a truly machine learning approach has not been successfully applied to the segmentation of the LCN. Thus far, researchers have relied on thresholding or region-growing to segment the LCN, which often require manual intervention to correct artefacts and remove background, sometimes pixel-by-pixel. Several approaches can also only handle one set of scans at a time, or one region at a time, and are not suitable to analyse a large portion of the osteocyte network. Kerschnitzki et al. employed a low-pass filter and adaptive thresholding to segregate the LCN^37^, though this approach is susceptible to image artefacts and capturing objects of larger size (e.g. blood vessels). Building on this to analyse connectomics of the LCN, Mabilleau et al. employed a thinning algorithm to refine the dendrites into a singular-voxel-wide skeleton^38^, and then computed the length and degree distributions of the network. Further application of this technique required meticulous manual examination to detect and manually draw in overlooked osteocytes and dendrites^39^. Similar semi-manual approaches were applied by other researchers, attempting to detect and segment osteocytes by fitting ellipsoids to cell bodies^40,41^. Additional work by Heveran et al. also incorporated ellipsoid shape priors and thresholding to develop an automatic segmentation and analysis technique of osteocyte lacunae within 3D CLSM images^25^. The model demonstrated notable predictive efficacy, achieving accurate segmentation of 77.1–97.8% of the lacunae in their experimental dataset^25^. Nevertheless, akin to other thresholding methodologies, their image pre-processing procedures necessitate manual adjustment of filtering and thresholding parameters when applied to different datasets.

A more complex recent automated segmentation method, variational region growing (VRG)^42^, has also been applied to the LCN, however it is initially limited by segmenting osteocytes and dendrites into the same class^43^. A number of additional studies attempt to combine with other methods to overcome this, but the method is highly sensitive to data availability and image resolution. Therefore, despite some early attempts, computer vision and machine learning methodologies have yet to be fully applied to efficiently measure the intricate architecture of the osteocyte network.

Given the biological complexity and computational challenges described, we first compare a range of automated thresholding methods with those applied previously, with the aim of identifying the most successful. Next, we train several CNN and transformer models on confocal osteocyte scans, improving and challenging their accuracy until a successful model is identified. In order to demonstrate the scientific and clinical potential of these techniques, we test the hypothesis that the model can distinguish between young and aged bone. Finally, this model is challenged to distinguish bone from genetically modified mice known to have LCN degeneration from their wild-type controls.

## Materials and Methods

### Fluorescent Imaging of Mouse Osteocyte Networks

Femurs for cryosectioning and fluorescent imaging were prepared as previously described^21,24,44,45^. Staining of the LCN was accomplished utilising the hydrophobic lipophilic dye, DiI (ThermoFisher), Alexa Fluor 488-Phalloidin (ThermoFisher), and DAPI for identification of cell nuclei. Bone sections were optically cleared and imaged using confocal laser scanning microscopy (CLSM) on a Leica DMi8 (Leica Microsystems) inverted microscope running LASX software.

Single femur bones from inbred, male C57BL/6 mice (young, 2 months, N=4 and aged, 36 months, N=5) from the Buck Institute for Research on Aging (Novato, CA) were used for comparisons regarding age-related phenotypes, with genetic modification-induced degeneration measured in femurs from male (young, 2 months) TβRII^ocy-/-^ mice and their TβRII^ctrl^ littermates (N=4-5 ea.)^21^.

While z-stacks of multiple LCNs in 3D were available, these were not applied in this study due to computational cost and high error rates in initial trials. Instead, individual scans of the osteocyte network were collected across multiple randomly selected regions in the femur, resulting in approximately 100-135 scans per group for testing the model.

### Datasets

Although the initial collection consisted of 1,224 images, constraints in labeling reduced the number of images suitable for training (892 images). In this study, we utilised two distinct datasets for initial model training and selection, followed by further training of the most successful model on an enlarged dataset.

*Dataset 1:* This comprises a manually annotated set with 30 training and 7 validation images. The manual labelling was facilitated by the automated image analysis tool, arivis Cloud (Carl Zeiss Microscopy GmbH, 2023).
*Dataset 2:* This dataset comprises 696 training images alongside 196 images designated for validation. The labels within this dataset are derived from a segmentation model previously trained on Dataset 1. As a result, these labels inherently possess certain biases and are primarily employed for training with partial labels. The segmentation model’s automatic training was facilitated using the arivis Cloud software.
*Dataset 3:* Finally, an additional 49 scans were annotated manually in order to further train the Attention U-Net. This required manual annotation by one trained and one expert operator, taking a total of 120 hours.

### Evaluation Metrics

In order to quantitatively evaluate the performance of the segmentation methods, Dice score coefficient (DSC) and Jaccard index or Intersect over Union (IoU) were employed. Both DSC and IoU are among the most commonly used metrics in semantic segmentation^27,46,47^. The equations are shown in (1) and (2) respectively where *P* - represents Prediction and *G* - Ground Truth. Dice metric evaluates the spatial overlap accuracy between the predicted segmentation and the ground truth. The Jaccard index (or IoU) is defined as the area of the intersection between the prediction and ground truth, divided by the area of the union of two label sets^46^.

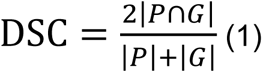

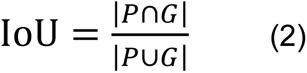

### Segmentation using thresholding

We replicated the segmentation methods described in Kerschnitzki (2013); Mabilleau et al.^38^ and in Ashique et al.^39^ (which we have termed M1, and M2 respectively), since they were also based on CLSM images. Owing to considerable variability in our data, such as bone sample preparation, spatial resolutions, dyeing agents etc. we optimised method parameters and threshold functions to our dataset. In addition to this, inspired by previous literature, we derived two additional threshold-based segmentation techniques. While both methodologies shared the same pre- and post-processing stages, they differed in their core segmentation techniques: one employed Canny edge detection for canaliculi segmentation, while the other opted for Otsu thresholding. The pre-processing began with a SciPy’s Gaussian filter with a standard deviation of 2 to decrease noise such that derivative-based thresholding functions were better facilitated. Secondly, we manually fit a binary threshold of 70 to separate the image into two masks, the lower end being osteocytes and the higher threshold being dendrites. We performed a second binary threshold on the dendrite mask to remove additional noise. Core segmentation techniques were then applied (Canny(p1=70, p2=220) and Otsu(p1=80, p1=255)). The post-processing employed a morphological closing operation on the dendrite mask using a 3×3 kernel. This facilitated the refinement of segmented dendrites. Lastly, overlapping osteocyte and dendrite labels were removed using an elementwise subtraction yielding the final segmentation.

### Segmentation using deep learning models

In the realm of medical image segmentation, certain architectures have risen to prominence due to their superior performance on a myriad of tasks. Particularly, U-Nets and Vision Transformers have recently been recognised for their state-of-the-art segmentation capabilities. Guided by this knowledge and the prevailing literature, our experiments primarily centred on leveraging these architectures for our specific segmentation requirements.

#### Image preprocessing and augmentation

Given that our RGB images only held information in the red channel, they were converted to grayscale to retain essential data while simplifying the input. To bolster data diversity and counteract potential overfitting of the models, various data augmentations, such as random flips, rotations, and cropping, were applied. Additionally, gradient norm clipping was incorporated, and datasets underwent normalisation before training.

#### Training parameters and strategy

Initial hyperparameters were retrieved from the original publications associated with each model. However, to ensure optimal performance on our specific datasets, these parameters underwent meticulous testing and subsequent adjustments. For our best performing models, we applied PyTorch’s implementation of the AdamW optimiser with learning rate = 0.001, weight decay = 1e–4 and cosine annealing scheduler with parameters T max = 10% of total steps and eta_min = 5e−6. For the loss function, we used a weighted sum of Dice loss and Cross Entropy loss: DiceCELoss. The mentioned preprocessing techniques and training parameters were used for all experimental training in this project. Training was conducted using the Andrena HPC facility at Queen Mary University of London. The computational tasks were executed on a single Nvidia A100 GPU.

#### Comparison of Dataset 1 and Dataset 2

The first experiment adopts the following architectures: U-Net, UNet++, RegU-Net, Flex U-Net, SwinUNet, and UNETR. These models were sourced from Medical Open Network for AI (MONAI), a PyTorch-based library tailored for medical imaging applications. These models were trained using two different training strategies: Strategy 1 where models were exclusively trained on Dataset 1, and Strategy 2 where initial training was conducted on Dataset 2, followed by fine-tuning on Dataset 1. The second strategy was implemented due to the time-consuming and challenging task of annotating 2D images from the original 3D stack of CLSM images (Fig.1). Notably, during the training on Dataset 1 in Strategy 2, the labels are partially annotated, hence, the loss function was exclusively computed over the annotated areas. As a result of this, we could leverage the segmentations predicted by a model trained on arivis Cloud on Dataset 1, thus giving access to a greater number of labels. The motivation behind this was to explore if increasing the dataset with sub-optimal labels could provide a better result compared with a much smaller training dataset but with better labels. The fine-tuning process in Strategy 2 was evaluated through three distinct strategies:

- Freezing all layers except input and classification layer
- Freezing all layers except classification layer
- Implementing a reduced learning rate on all layers, except the classification layer

#### Separating model prediction

The model aimed to utilise mutual information and contextual cues from both osteocytes and dendrites in the image to learn intricate features. For instance, the gathering of numerous dendrites within a specific region could signify the presence of an osteocyte. Capturing such nuanced and interrelated features might necessitate a more extensive annotated dataset than was available to us. In light of these considerations, we also embarked on an alternative strategy: training two distinct models—one focusing on dendrite identification, and the other on osteocyte prediction. The aggregated outputs from these models yielded the final segmentation.

#### Encouraging longer and fewer dendrites

Moreover, in order to leverage the models’ segmentation outputs for subsequent connectomics analysis, it is also imperative that the model is able to predict dendrites that establish clear interconnections between osteocytes. In order to encourage the model to output longer and interconnected dendrites, we incorporate an additional means of regularisation on the loss function as seen in Algorithm 1. This was done by first performing a connected components analysis of the dendrite prediction. We then calculated the size of each component and added an inverse penalty; smaller components led to larger penalties. The motivation behind this strategy was essentially to penalise the network for outputting short, isolated dendrites. Two specific models were employed for this endeavour: the CNN-based Attention U-Net and the transformer network Swin UNetR.

### Connectomics analysis

To facilitate a quantitative connectomics analysis of the segmentation outputs, the outputs were transformed into a graph representation using the Python library NetworkX. In this structured representation, osteocytes serve as nodes, with dendrites forming the interlinking edges between them. To achieve this, the segmentations were initially separated into two binary masks: Osteocytes and Dendrites. Subsequently, a morphological dilation was applied to the masks using a 4×4 ellipsoidal structure for Osteocytes and a 2×2 cross-structuring element for Dendrites. This dilation procedure aimed to merge nearby components like noisy dendrite predictions that, ideally, should represent a singular entity. After dilation, a connected component analysis was executed on both classes to individually discern unique components. To establish edges that link osteocyte nodes, an iterative process was undertaken on dendrite segments, verifying overlap with osteocyte labels based on their x and y coordinates. This framework further facilitated the computation of metrics, encompassing the network’s degree, the shortest path between osteocytes, the frequency of terminus connections per osteocyte (dead-ends), and dendrite thickness across the network and individual connections. These metrics were manually tested and validated on simplified dummy data.

### Statistical Analysis

In order to test the success of the model at distinguishing between different experimental groups, statistical analyses performed using a two-sided student’s t-test with a defined alpha level of 0.0001 (indicated by **** on figures). Prism 8.4 (GraphPad Software Inc.) was used for all statistical comparisons. Unless otherwise stated, data is presented as mean ± SD, and groups are detailed in the figure legends.

## Results

### Improved thresholding approaches outperform previous methods for segmenting the osteocyte network

Average segmentation time, Dice score, mean IoU (mIoU) and IoU per class is presented in Table 1 for each of the thresholding methods uses. M1 is based on previous thresholding methods of Kerschnitzki et al.^37^, while M2 replicated the manual adjustments to thresholding segmentation applied by Mabilleau et al.^38^ and in Ashique et al.^39^

**TABLE 1:**
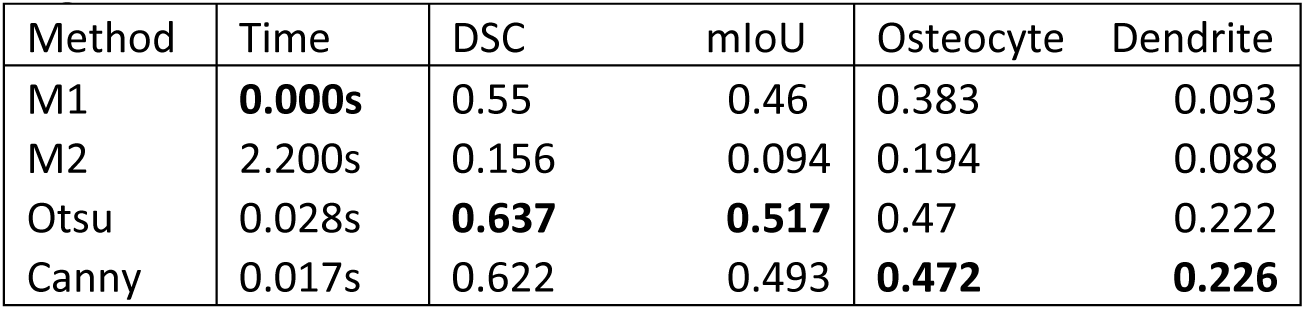
Segmentation time, Dice score (DSC), mean IoU and IoU for individual classes. The methods M1 and M2 are derived from (Kerschnitzki, 2013; Mabilleau et al., 2016)^37,38^, and (Ashique et al., 2017)^39^ respectively, Otsu and Canny represent our non-DL segmentation methods.

When comparing these previously applied methods with Otsu and Canny approaches we developed, we found that our methods significantly outperformed them. While both of our methods thresholded osteocytes and dendrites with similar accuracies (∼47% and ∼22%, or 0.47 and 0.22 mIoU, respectively), the Otsu method gave the highest confidence, with a Dice score of 0.637 and thus was the most promising segmentation methods among the four thresholding approaches. While M1 had an advantage in execution speed, its segmentation performance lagged behind Otsu and Canny. M2 performed significantly worse in all metrics compared to the other methods.

### Attention U-Net is the most effective deep learning model capable of segmenting osteocytes and their dendritic processes

Comparison of training strategy 1 and 2: performance metrics of the models from the two different training strategies can be viewed in Table 2. Among the evaluated models, the Attention U-Net achieves the highest Dice score and mIoU for both strategies. In contrast, the RegU-Net model displays notably lower performance than the other models in both the Dice score and mIoU. The performance of UNet and U-Net++ are quite similar for training strategy 1, and inconsistencies arise in strategy 2. The performances of the Vision Transformers (UNETR and Swin UNETR) are relatively lower than the top models but are still respectable. Swin UNETR’s performance is notably inconsistent between the training strategies. It performs well on Dataset 1 (only slightly below U-Net), but its performance drops significantly in strategy 2.

**TABLE 2:** Performance metrics of implemented models on two different training strategies.

|  | Training strategy 1 (Dataset 1) |  |  |  | Training strategy 2 (Dataset 2 + fine-tune on Dataset 1) |  |  |  |
| --- | --- | --- | --- | --- | --- | --- | --- | --- |
| Models | DSC | mIoU | Osteocyte | Dendrite | DSC | mIoU | Osteocyte | Dendrite |
| U-Net | 0.795 | 0.694 | 0.773 | 0.389 | 0.195 | 0.117 | 0.015 | 0.088 |
| U-Net++ | 0.794 | 0.694 | 0.779 | 0.385 | 0.794 | 0.694 | 0.78 | 0.383 |
| Attention U-Net | **0.801** | **0.703** | **0.793** | **0.394** | **0.8** | **0.702** | **0.794** | **0.391** |
| RegU-Net | 0.523 | 0.432 | 0.393 | 0.05 | 0.504 | 0.431 | 0.402 | 0.005 |
| UNETR | 0.76 | 0.659 | 0.755 | 0.301 | 0.759 | 0.658 | 0.755 | 0.305 |
| Swin UNETR | 0.79 | 0.69 | 0.782 | 0.37 | 0.234 | 0.36 | 0.113 | 0.179 |

#### Experiments to improve dendrite segmentation accuracy

The outcomes of the three experiments—namely, separating model predictions, implementing a novel regularization technique to promote larger yet fewer dendrites, and employing label dilation to account for annotation inaccuracies—are summarised in Table 3. The findings demonstrate that the initial two experiments exert minimal impact on the segmentation accuracy of the dendrite class. In contrast, the approach involving the dilation of dendrite labels prior to training yields a notable enhancement in dendrite accuracy, registering an increase of 25% for the Swin UNet and 22% for the Attention U-Net. Among the models examined, the Attention UNet consistently exhibits superior performance. Superficially, dendrite predictions appear satisfactory when studying the segmentation results depicted in Fig. 2. However, upon closer inspection, we can observe the presence of fragmented and disconnected dendrite structures.

**Fig. 2:**
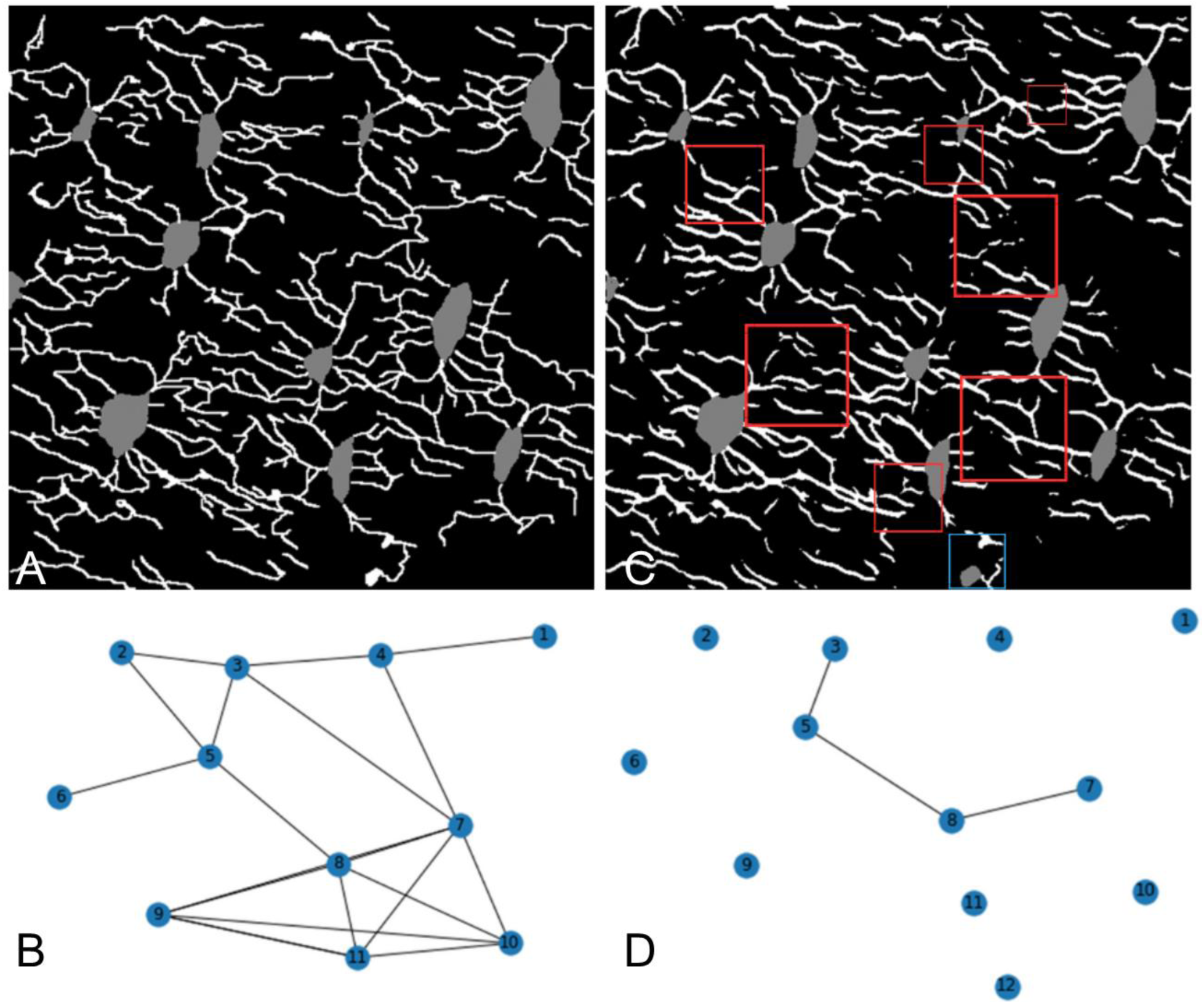
Connectomics used as training strategy for deep learning models. (A) Annotated labels in an image, and (B) the corresponding connectomics map. After training, model is tested to produce a (C) prediction and (D) corresponding connectomics map, which can be compared to the annotated map to assess accuracy. Red squares in the prediction highlight instances of disconnected dendrite segments, while the blue square indicates a misclassification where a dendrite has been erroneously labeled as an osteocyte.

**TABLE 3:**
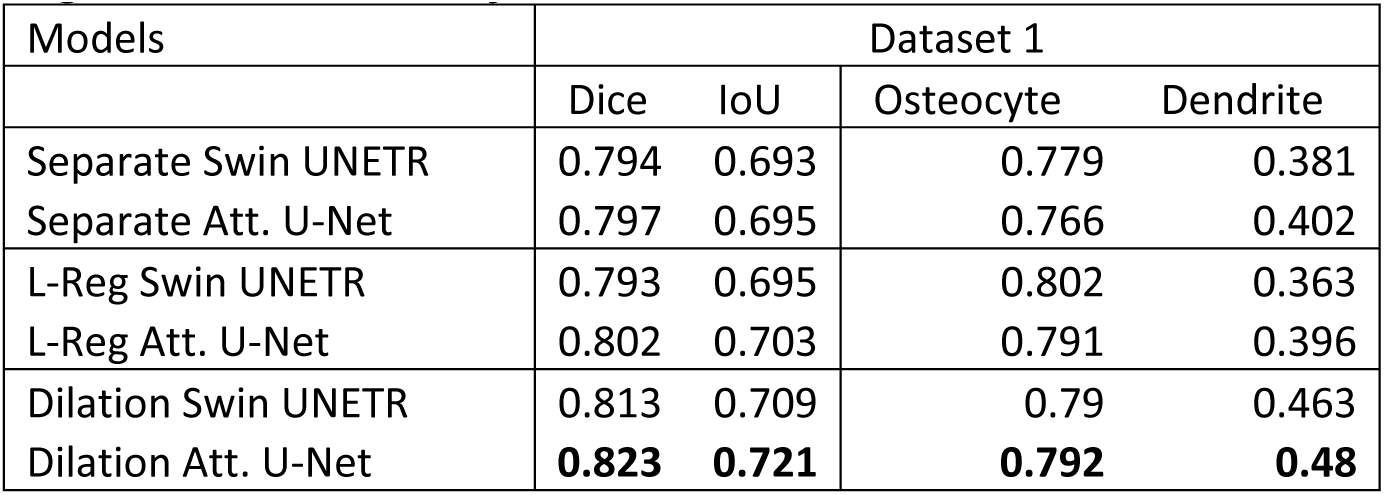
Performance metrics of experiments attempting to improve dendrite segmentation accuracy.

Connectomics analysis: The connectomics analysis in Table 4 was conducted using segmentation predictions from the top-performing model highlighted in Table III. This model exhibits a tendency to overestimate the number of nodes, predicting an average of 8.30 nodes as opposed to the 7.00 nodes present in the labelled data. Notably, the model’s predictions present fewer dead ends per node, averaging at 14.9, in contrast to the labelled data’s 20.9. Furthermore, it significantly undervalues the average number of connections per node, registering a mere 0.41 compared to the 3.84 seen in the labels. From the analysis, we can also see that the model produces shorter and thicker connections than the labels. These findings are also reflected in the comparison of a label’s graph and a segmentation’s graph in Fig. 2.

**TABLE 4:**
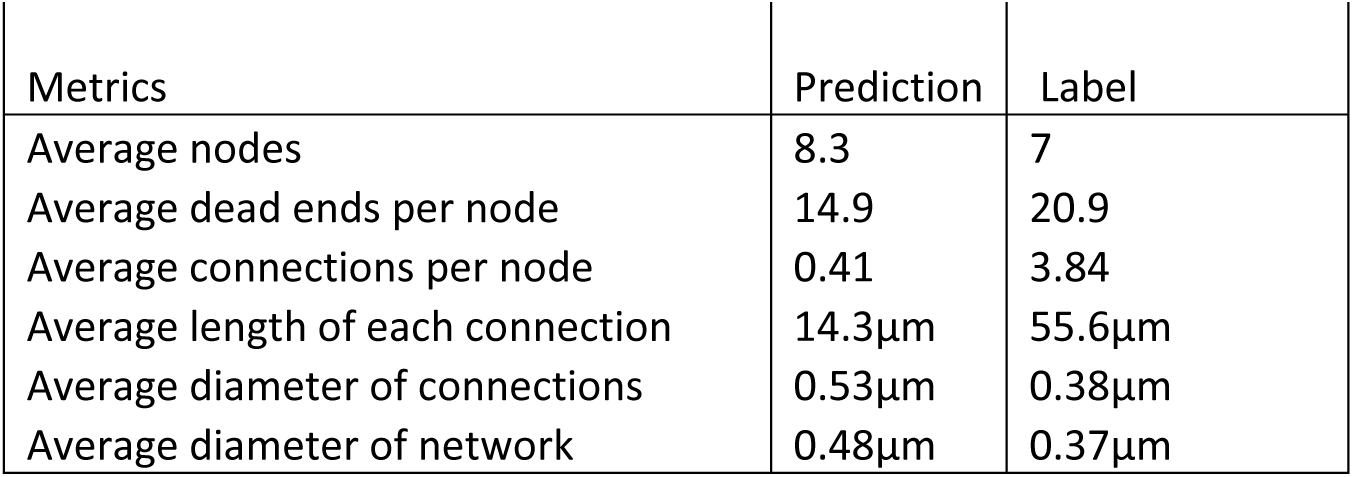
Connectivity metrics computed from the connectomics analysis. The average is taken from the validation set consisting of 7 images.

Having now selected the Attention U-Net as the most successful model, an additional 49 images were manually annotated to increase the training dataset (Dataset 3). To demonstrate the labour-intensive nature of this manual segmentation, pixel-by-pixel labelling of these datasets by an expert operator took more than 120 hours. Upon re-training the model, the accuracy of osteocyte detection increases to 81.8%, while the accuracy of dendrite detection increased to 41.2% when compared to an expert operator. While the accuracy of osteocyte detection was very high, it remained unclear whether the lower dendrite accuracy would be sufficiently high to distinguish between two populations, e.g. young and aged bone.

### Deep learning models can successfully distinguish between young and aged, or healthy and degenerated, osteocyte networks without manual intervention

Work by the authors has previously demonstrated that aging disrupts the integrity and predicted function of the osteocyte network, using labour-intensive computational modelling techniques to manually segment and monitor these changes, and relating them to age-related changes in gene expression^21^. Given that we have identified several parameters of the osteocyte network that are quantifiably different in aging when compared to younger mice, we challenged our deep learning model to similarly distinguish between the two.

We found that the model was capable of correctly identifying and measuring the reduced connectivity of the network in 2D confocal microscopy scans of older mice, and the decreased average length of dendritic connections (Fig.3). It also correctly found a reduced number of osteocytes in the aged bone, as well as half the number of connections per osteocyte, corroborating our manual measurements in our previous work. While an additional measure of average dendrite thickness was measured, and no difference was found between young and aged mice, this measure may be much more sensitive to user error and image quality and so is significantly less reliable.

**Fig. 3:**
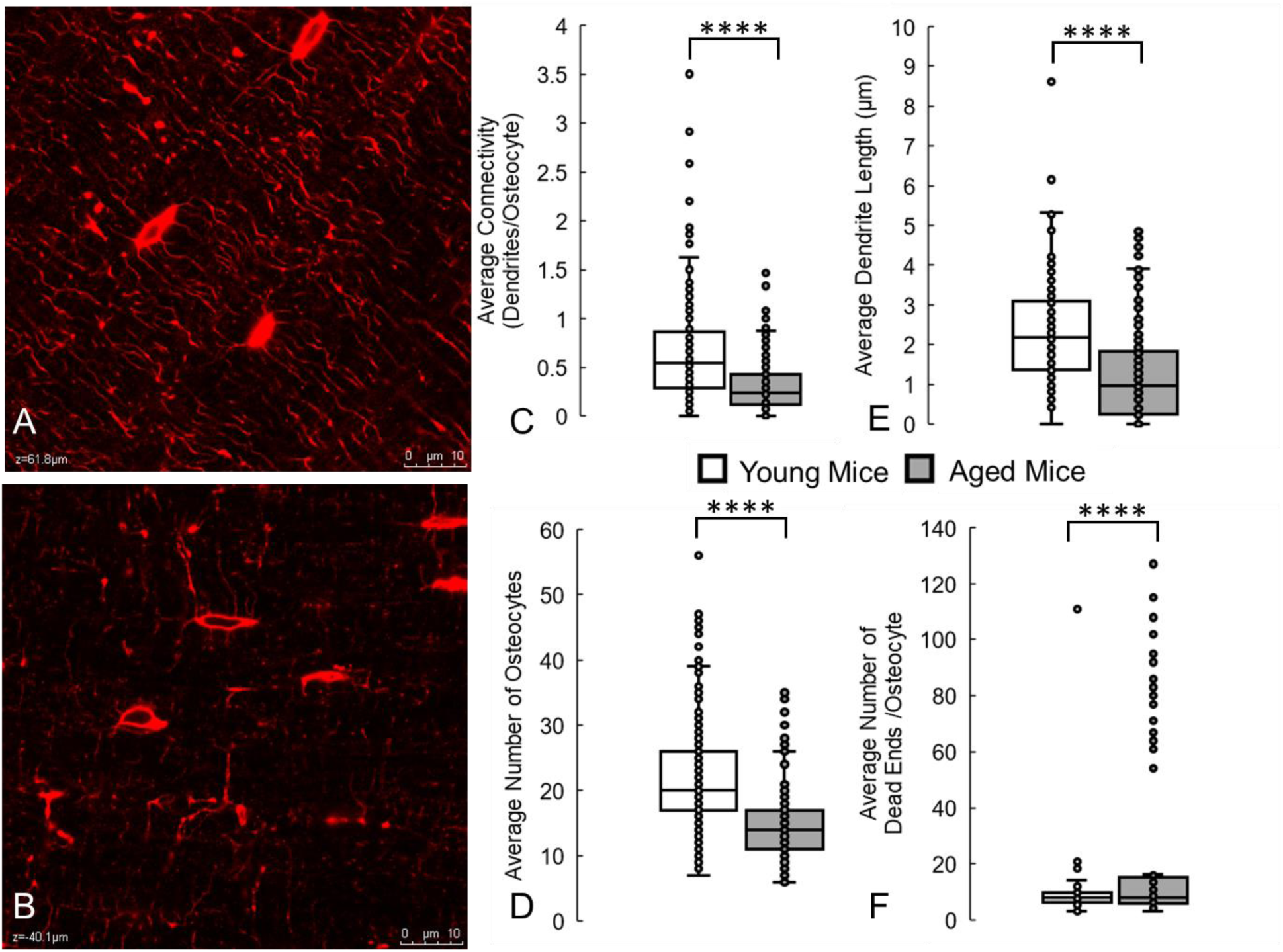
Deep learning model can accurately detect and identify the lower osteocyte connectivity of aged bone, when compared to young bone. These results are similar to changes we measured manually in a previous study^21^. (A) Confocal laser scanning microscopy images of osteocytes (red, phalloidin, cytoskeleton) from young (2 month old) and (B) aged (36 month old) mice. Running the trained deep learning model on these datasets correctly measured that aging causes significant decreases in (C) average network connectivity, (D) number of osteocytes, and (E) average dendrite length. (F) The model also correctly predicted significant increases in blunted dendrites or dead ends in the network. **** denotes *p* < 0.0001 by Student’s t test compared to young control. Scale bar = 10 µm.

Similarly, our deep learning model could partially predict the degeneration in the LCN induced by ablation of TGF-β Receptor II in osteocytes (Fig.4), capturing the trends of decreased connectivity and greater blunted canaliculi that we measured manual in our previous study^21^. However, the model did not correctly capture the decreases in number of osteocytes and dendrite length that we found previously. This may indicate that training on larger datasets of LCNs from healthy bone is required to pick up the more subtle degeneration driven by individual genetic mutations compared to the broad degeneration generated by aging.

**Fig. 4:**
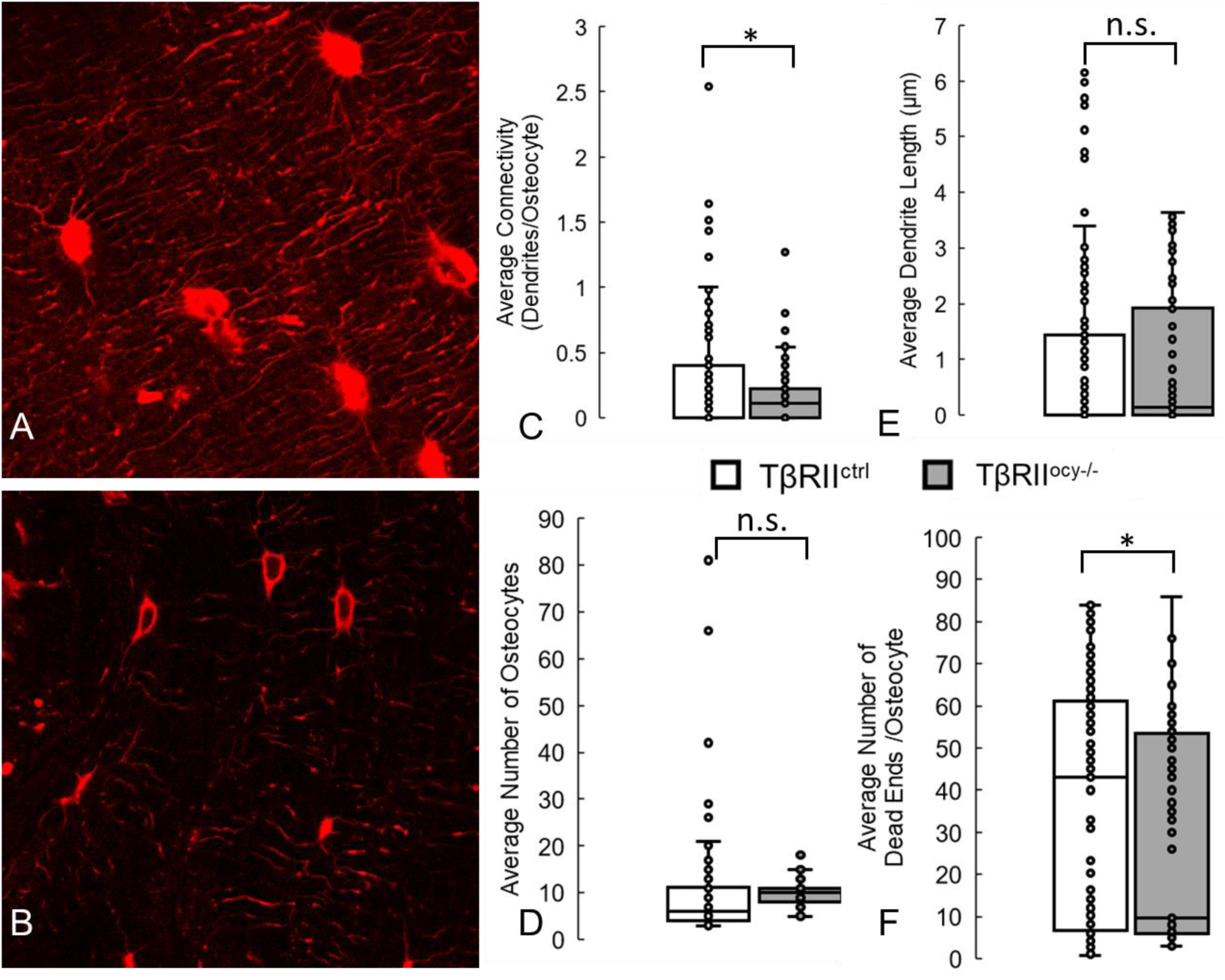
Deep learning model can detect and identify the lower osteocyte connectivity of bones in which TGF-β Receptor II has been ablated from osteocytes, compared with control mice. Some of these results (decreased connectivity and number of dead ends) are similar to those measured by us previously, while others (decreased osteocytes and dendrite length) are not fully captured^21^. (A) Confocal laser scanning microscopy images of osteocytes (red, phalloidin, cytoskeleton) from 2 month old control mice (TβRII^ctrl^) and (B) their litter-mates in which TGF-β Receptor II was ablated from osteocytes (TβRII^ocy-/-^). Running the trained deep learning model on these datasets correctly measured that aging causes significant decreases in average network connectivity and average number of blunted canaliculi in the network, but not previously observed decreases in number of osteocytes and average dendrite length. * denotes *p* < 0.05 by Student’s t test compared to control, n.s. denotes no statistical significance. Scale bar = 10 µm.

Taken together, these results demonstrated the capacity of deep learning models, and the Attention U-Net in particular, to automatically characterise the age and integrity of the osteocyte network, providing a powerful new tool to analyse bone structure at the cellular scale.

## Discussion

This study presents, for the first time, a deep learning model capable of automatically segmenting the osteocyte network from microscopy scans. Testing a range of deep learning techniques, we identify Otsu as an effective thresholding method, and an Attention U-net as the most effective convolutional neural network for this task. In doing so, our model is capable of segmenting osteocytes and dendrites with 81.8% and 41.2% accuracy, respectively. Importantly, this model successfully distinguishes between the osteocyte networks of young and aged bone, and degeneration induced by genetic modifications, matching our previously published results and validating the model.

In taking on such a challenging computational task, a number of limitations were necessary. Primarily, the restricted size of our dataset appears to be a pivotal constraint, leading to the model’s plateauing performance and resulting in fragmented dendrite segmentations and occasional misclassifications of osteocytes. The limited dataset also affects our quantitative assessments of LCN connectivity. Moreover, there is an absence of a benchmark dataset on the topic, making it challenging to compare our segmentation model or connectomics analysis methodology with prior endeavours outside our own previous works. To enhance our outcomes, expanding the dataset is an evident avenue of further enquiry. Furthermore, while the manual annotation tool we utilised supported the resolution of our original images, the actual resolution proved somewhat inadequate for the precise annotation of thin structures. The dendrites typically had a pixel thickness ranging from 1.5 to 3 pixels, making their annotation exceptionally meticulous and challenging. Consequently, even minor discrepancies in annotations could lead to considerable variances in results. While utmost care was exerted to ensure the highest possible accuracy, including manual annotations by an expert operator with >15 years of experience of osteocyte segmentation, inter-operator variability may affect results and the practical challenges of zooming in and out, combined with the dendrites’ innate thinness, meant that some degree of imprecision was unavoidable. Furthermore, additional future research could study inter-observer error, which could be high for the dendrites. To mitigate the potential inconsistencies arising from this, we implemented an experimental strategy: a morphological dilation of the dendrite labels to slightly enhance their thickness (2×2, and 3×3 kernels). By doing so, we aimed to ensure a more accurate encapsulation of the dendrites as observed in the original images. Given the inherently slender nature of dendrites, we postulated that it would be beneficial to slightly overestimate their thickness in the labels rather than underestimate it. However, this is a limitation that should be addressed as models are further developed in future.

It is interesting that in using a relatively simple comparison of thresholding methods, we have found that a well-known thresholding operation in biomedical image analysis, Otsu, was both a simpler and more accurate method for our dataset than those applied previously. This is because these thresholding approaches need to manually adapted to each dataset. In particular, an accepted laborious task in the study of osteocytes is the manual segmentation of all of these individual cell bodies and dendrites for the study of their biochemistry, biomechanics, mechanobiology, etc. As previous methods had a significant manual component, the thresholding method we have studied could significantly simplify the preparation of osteocyte CLSM scans for many researchers. The further development of these techniques to automate the segmentation process, even at the current accuracy, would massively reduce what is currently an extremely labour-intensive task that relies on expert operators with years of training.

A particular challenge for these models appears to be accurate segmentation of dendrites, with our model achieving less than 50%. This may occur due to the convoluted trajectories of individual dendrites and canaliculi through the bone matrix in 3D, meaning that a confocal plane is highly unlikely to capture the path in its entirety. However, as AI models usually improve exponentially with additional data, and only a few hundred 2D scans were used in this study, this could likely be significantly improved with additional large datasets. Furthermore, with larger datasets and matched resources for segmentation, this analysis could be extended to 3D across z-stacks in order to capture changes in the LCN in its entirety. Another notable result is the lack of change measured in dendrite or canalicular thickness, because if this parameter could be measured, then it would provide a significant boost to distinguish spatial differences in remodeling around osteocyte cell bodies or dendrites in the challenging new research area of peri-lacunocanalicular remodelling (PLR). This insensitivity to thickness changes is already a challenge due to the limitations of the wavelength of light, which is generally larger than the thickness of these changes. However, this is a challenge to which self-supervised machine learning may be well suited, as this type of pattern recognition is much better suited to computer vision. This holds forth the tantalising prospect that future work could include self-supervised learning methods to enable us to actually measure the changes in canalicular and lacunar width associated with PLR.

Most excitingly, this first attempt to apply AI and computer vision to map the osteocyte network is already capable of distinguishing between the connectomics of young and aged bone to a very high degree of confidence. Additionally, it could measure a number of degenerative patterns induced by genetic modifications known to disrupt the osteocyte network, indicating that additional training could further improve this model as a tool for osteocyte analysis. All of the parameters measured matched those previously quantified by our team using a manual image processing and connectomics approach^21^, and thus this vastly improves the speed at which osteocyte network integrity can be quantified. Therefore, in summary, our deep learning model represents a powerful new research tool, of immediate value to the researchers in multiple fields of bone research.

## Acknowledgements

This work has been funded by the European Union’s Horizon 2020 research and innovation programme under the Marie Sklodowska-Curie grant agreement No. 748305 (SWV). This research was supported by NIH National Institute of Dental and Craniofacial Research Grant R01 DE019284 (T.A.) and NIH National Institute on Aging Grant F31 AG063402 (C.A.S.). UCSF cores used to complete this work include the Skeletal Biology and Biomechanics Core of the Core Center for Musculoskeletal Biology and Medicine (NIH National Institute of Arthritis and Musculoskeletal and Skin Diseases Grant P30 AR066262) and the Biological Imaging and Development Core and staff for support using Imaris. This work forms part of the research portfolio of the National Institute for Health Research Barts Biomedical Research Centre (NIHR203330).

## Author contributions

Conceptualization: GGS, SWV,

Methodology: SDV, CAS, GGS

Investigation: SDV

Visualization: SDV, CAS, SWV

Supervision: TA, GGS, SWV

Writing—original draft: SDV, SWV

Writing—review & editing: SDV, CAS, TA, GGS, SWV

## Data and materials availability

All data are available in the main text or the supplementary materials.

## Algorithm 1 Regularisation Loss for Dendrites

Pseudocode of algorithm used to incorporate additional means of regularisation on the loss function

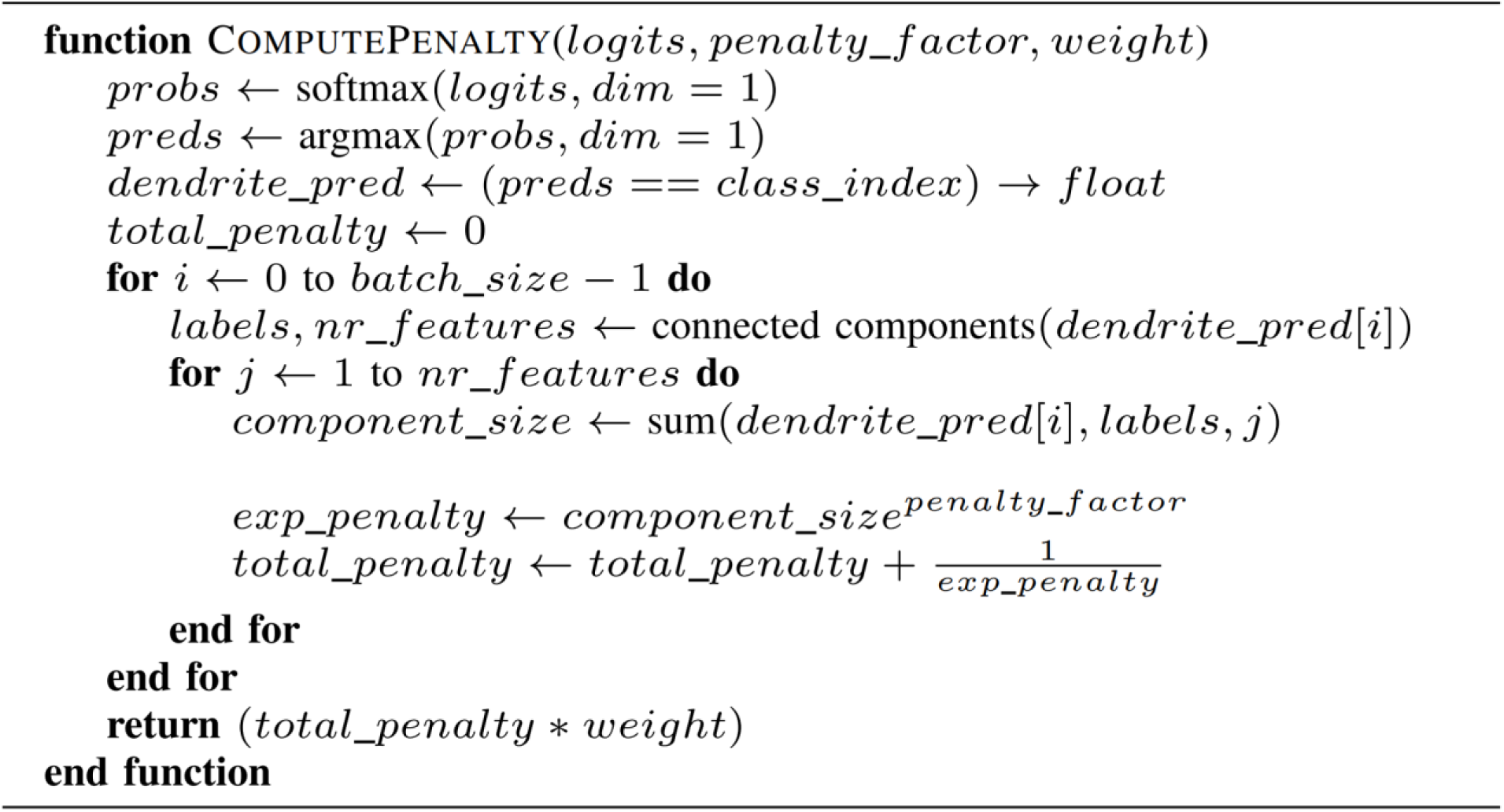

